# Consolidation of human skill linked to waking hippocampo-neocortical replay

**DOI:** 10.1101/2021.04.07.438819

**Authors:** ER Buch, L Claudino, R Quentin, M Bönstrup, LG Cohen

**Affiliations:** Human Cortical Physiology and Neurorehabilitation Section, NINDS, NIH, Bethesda, MD, USA

**Keywords:** replay, human motor learning, skill learning, offline learning, consolidation, generalization, hippocampus, entorhinal cortex, sensorimotor cortex, MEG

## Abstract

The introduction of rest intervals interspersed with practice strengthens wakeful consolidation of skill. The mechanisms by which the brain binds discrete action representations into consolidated, highly temporally-resolved skill sequences during waking rest are not known. To address this question, we recorded magnetoencephalography (MEG) during acquisition and rapid consolidation of a sequential motor skill. We report the presence of highly prominent, fast waking neural replay during the same rest periods in which rapid consolidation occurs. The observed replay was temporally compressed by approximately 20x relative to the acquired skill, occurred in both forward and reverse directions, was selective for the trained sequence and predicted the magnitude of skill consolidation. Replay representations extended beyond the hippocampus and entrorhinal cortex to the contralateral sensorimotor cortex. These results document the presence of robust hippocampo-neocortical replay supporting rapid wakeful consolidation of skill.

## Introduction

The popular idiom, *“practice makes perfect”*, emphasizes the importance of intense practice in acquiring and perfecting skill^1^. However, the same amount of practice results in significantly different skill depending on the presence or absence of waking rest within the training schedule, a phenomena termed the spacing effect^2^. Consolidation of skill is superior when frequent rest periods are interspersed with practice blocks (distributed practice) than when the same total amount of practice is performed over longer continuous blocks (massed practice)^2, 3, 4, 5, 6^. Thus, the presence of waking rest interleaved with practice is a crucial determinant of skill memory consolidation.

Consistent with this notion, it has been proposed that “*much, if not all*” skill learning occurs offline during rest rather than during actual practice^7^. For example, performance improvements while acquiring a new skill accumulate almost exclusively during waking rest periods interleaved with practice^8^. These *micro-offline* gains indicate a rapid form of skill memory consolidation that develops over a much shorter timescale than previously thought^8, 9, 10^, and that is approximately four times greater in magnitude than classically studied overnight consolidation requiring sleep^8^.

Skill learning involves forming new memories through binding, a process that hierarchically links simple action elements (e.g. – a single piano key press) to construct representations of complex spatiotemporal sequences (e.g. – a refrain within a classical piano concerto)^11, 12, 13, 14^. How the brain binds sequences of discrete action representations into consolidated, temporally precise skills during waking rest is not known^2, 4, 9^. One possible candidate mechanism is neural replay, the temporally-compressed reactivation of neural activity patterns representing behavioral sequences during rest. Previous work in rodents has shown that hippocampal replay constructs representations of navigational maps through relational binding of discrete spatial locations into trajectories^15^. Hippocampal activity has been documented during rest periods of early learning^9, 16^. Thus, it is possible that waking replay, through offline recapitulation of prior practice^15, 17, 18, 19^, promotes wakeful consolidation of skill; an issue that has not been investigated in humans or animal models^15, 20^.

Here, we posited a contribution of neural replay in wakeful memory consolidation. We tested this by applying a multivariate analytical approach to magnetoencephalography (MEG) brain activity data acquired during skill learning to decode spontaneous, waking replay during the periods of rest in which sequences of simple discrete actions learned during interspersed practice are bound into a consolidated skill.

## Results

Wakeful consolidation of skill was evaluated using a keypress sequence task^13, 21^ that shows rapid memory strengthening over a single session^8^. Subjects were instructed to type the sequence, *41324*, on a keyboard as fast and accurately as possible during each 10-second practice period. Thirty-six 10-second practice periods (trials) were interleaved with 10-second rest periods (Fig. 1). Skill was measured as the correct sequence typing speed (sequences per second, seq/s). Early learning, encompassing the set of trials within which 95% of total learning occurred, was explained by *micro-offline*, but not *micro-online* gains (Fig. 1D). That is, skill improvement occurred during waking rest, but not during active practice (Fig. 1C and D).

**Figure 1.**
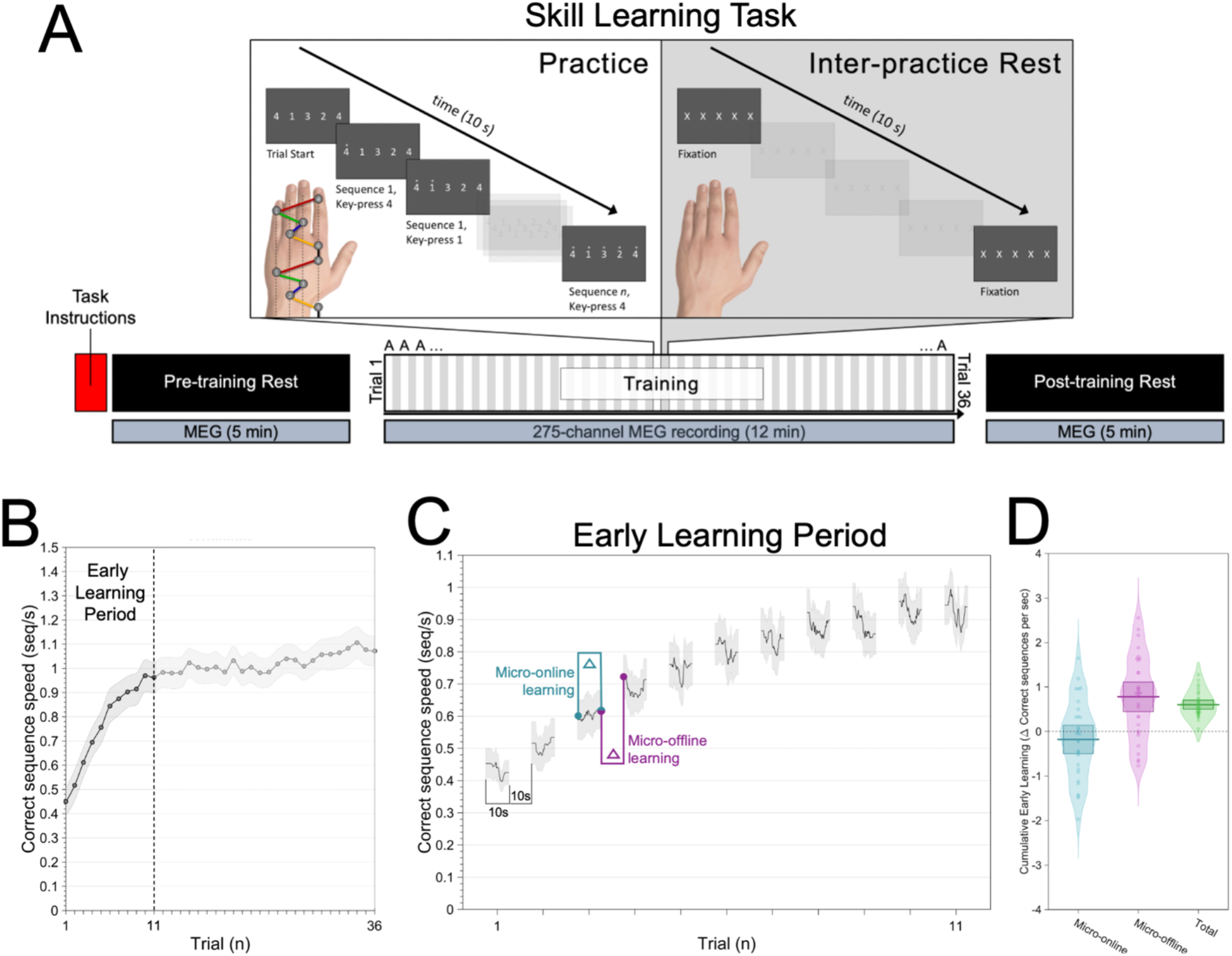
Skill learning task and behavioral performance. (A) Keypress sequence motor skill task. Subjects acquired a novel motor skill over a single training session. They were instructed to repeatedly type a trained sequence, *41324*, with the left non-dominant hand as fast and as accurately as possible. Keypress *4* was performed with the left index finger, Keypress *3* with the left middle finger, Keypress *2* with the left ring finger and Keypress *1* with the left little finger. Following task instructions, a 5-minute MEG waking rest recording was acquired prior to commencement of training (Pre-training rest). Subjects then performed the task over 36 individual practice periods lasting 10s each. Practice periods were interleaved with 10s waking rest intervals (Inter-practice rest). MEG was acquired continuously during training (12 minutes). A second, 5-minute waking rest MEG recording was acquired after training concluded (Post-training rest). (B) Performance curve. Skill was measured as the correct sequence typing speed (sequences per second, seq/s). Mean average performance (mean ± SEM) increased rapidly during early learning (the set of trials within which 95% of total learning occurred, trials 1-11; vertical dashed line). (C) Instantaneous correct sequence typing speed (group mean ± SEM shown) was used to quantify micro-online performance gains during practice (cyan) and micro-offline gains during interleaved rest (magenta) for the early learning period. (D) Micro-online (cyan), micro-offline (magenta), and total early learning (green) for each subject. Note that skill increases occur during intervening periods of waking rest, and not during active practice.

### Fast replay of skill sequences is prominent during waking rest interspersed with practice

Next, we examined parcellated, source-space MEG brain activity data for evidence of spontaneous replay of the trained sequence during waking rest (Fig. 2). We used Radial-basis Support Vector Machine decoders to interrogate resting-state MEG data for individual replay events (see Methods for details). Practice data segments centered on individual keypress events were used to train four one-versus-all keypress state decoders (i.e. – one decoder per finger). We then applied these decoders to waking rest MEG data to detect replay events (Fig. 2A) that occurred during pre-training, inter-practice and post-training rest periods. Since replay is known to be temporally compressed to different degrees relative to behavior, we looked for replay over a range of durations (25-2500ms) corresponding to different compression magnitudes (approximately 0.4-40x) relative to the acquired skill (the average correct sequence duration by the end of early learning was 1037.7±61.7ms; Mean±SEM; Fig. 1B).

**Figure 2.**
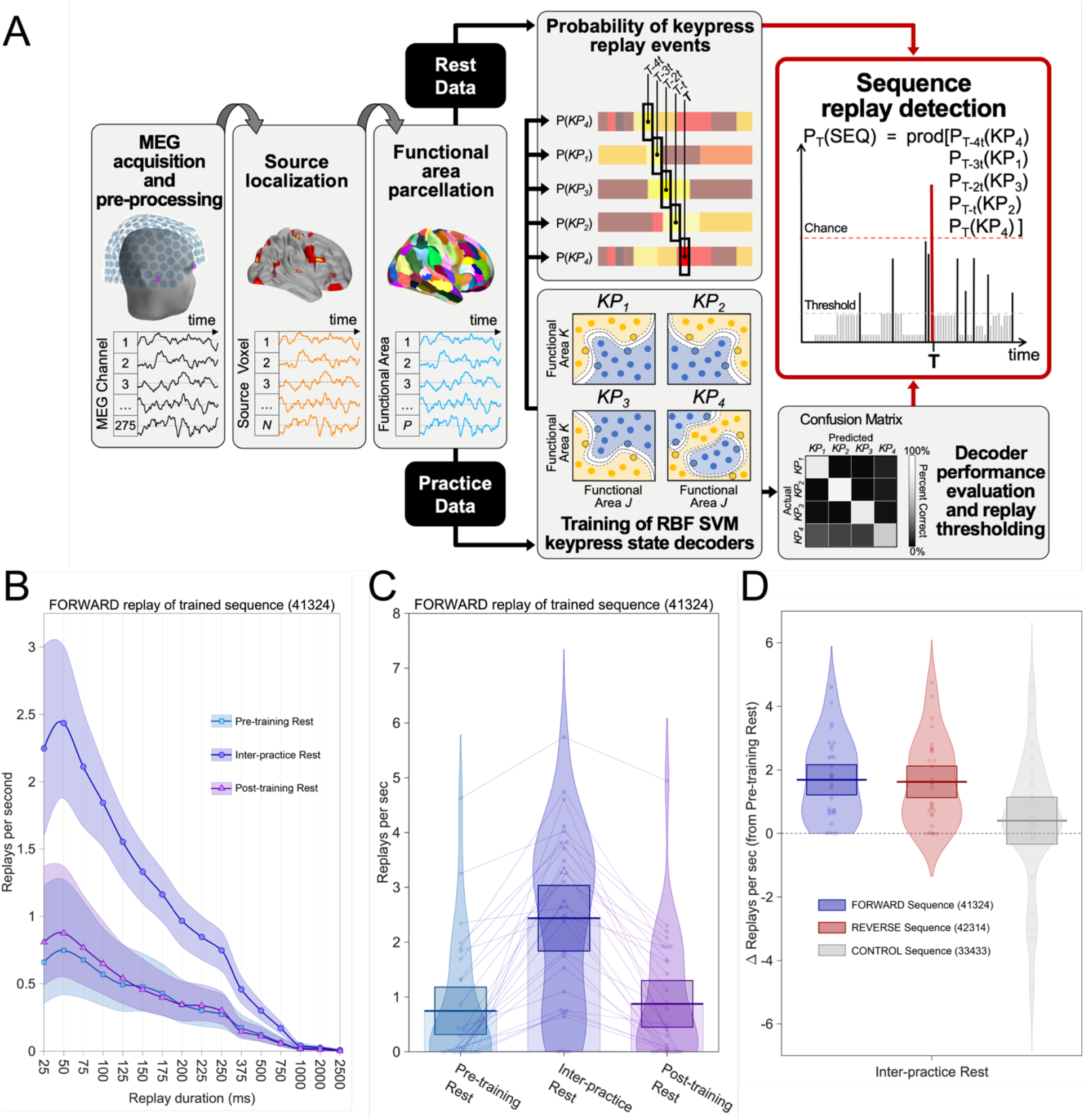
Replay detections. (A) Replay detection analysis pipeline. Raw MEG data was pre-processed, source-localized and parcellated into 216 brain regions using the Brainnetome Atlas^50^. Practice data segments centered on individual keypress events were used to train four one-versus-all Radial-basis Support Vector Machine keypress state decoders (i.e. – one decoder per finger). We then used the trained keypress state decoders to interrogate resting-state MEG data (preceding training, during rest periods interspersed with practice, and after the end of training) at each time-point, T, for replay of keypress state sequences over a range durations (25-2500ms) corresponding to temporal compression ranges (0.4-40x) previously reported^26, 27, 28^. Decoder performance evaluation (Fig. S1) was used to threshold minimum detectable sequence replay probabilities. Permutation testing against probabilities for all possible sequences (n=1024) at time-point T was used to determine statistically significant detection of keypress sequence replay events. (B) Neural replay was detected during waking rest. (A) The maximum forward trained sequence replay rate was observed for events of 50ms duration. This was consistent with observed replay for the reverse trained sequence (Fig. S2A). (C) Waking replay rates (50ms duration) for the forward trained sequence more than tripled during inter-practice rest (2.44±0.29; Mean±SEM) relative to pre-training rest (0.75±0.21). Similar magnitude rate changes were also observed for the reverse trained sequence (Fig. S2B). Replay rates then receded during post-training rest (0.88±0.21) to levels similar to pre-training. (D) Within individuals, waking replay increased more prominently (F_2,87_=6.42; p=0.002) after practice onset for the forward (*41324*; left; p=0.0062, Bonferroni-corrected) and reverse (*42314*; middle; p=0.0099, Bonferroni-corrected) trained sequence than for a control sequence (*33433*; right). This control was selected on the basis that it was the only one out of 1,022 possible alternate sequences sharing no common ordinal position or transition structure with the forward (*41324*) and reverse (*42314*) trained sequence. Pre-training rest replay rates did not significantly differ between these sequences (F_2,53.94_=2.431; p 0.098). Of note, this same trend was observed even after expanding this criteria to consider additional sequences sharing either a single common ordinal position or a single common transition with the practice sequence (Fig. S2C).

Neural replay of the full trained sequence was identified in all waking rest periods examined. However, detection rates (i.e. – the number of replay events detected per second) varied depending on replay duration (i.e. – compression magnitude; Fig. 2B). Forward replay rates for the trained sequence (*41324*) were most prominent for events lasting 50ms and progressively decreased for longer event durations (Fig. 2B; see Fig. S2A for reverse replay of the trained sequence, *42314*). Thus, the modal neural replay representation of the skill was temporally compressed approximately 20x relative to the actual behavior. Figure 2C highlights the large difference in forward replay rates across the different waking rest states (F_2_,58=15.26, p=0.000018). Specifically, forward replay events during inter-practice rest occurred approximately 3 times more frequently (2.44±0.29 replays/s) relative to pre-training (0.75±0.21 replays/s, p=0.00528) or post-training rest (0.88±0.21 replays/s, p=0.00089, Fig 2C), which were both comparable (p=0.2972). Reverse replay of the trained sequence exhibited similar features (Fig. S2B). Elevated replay rates were maintained across all individual inter-practice rest periods (Fig. S3A). Within each of these rest periods, replay events were likely to occur in clusters (Fig. S3B,D) that were uniformally distributed across the entire 10 seconds (Fig. S3B,C). Finally, mean replay rates during early learning (initial 11 trials) were significantly higher than during late learning (last 11 trials; t_29_ = 2.66, p = 0.006, one-tailed).

We also interrogated resting-state MEG data for fictive replay of a single unpracticed control sequence (*33433*). This sequence was selected on the basis that it is the only unpracticed sequence combination (out of 1,022 possible alternatives) that shares no common ordinal position or transition structure with forward and reverse replay of the practiced one. This analysis rendered two important results. First, we detected fictive replay of this unpracticed control sequence in all waking rest periods examined, and with comparable pretraining-rest rates to both forward and reverse replay of the practiced one. However, after practice onset there was a greater increase (F_2,87_=6.42; p=0.002) for both forward (p=0.0062; Bonferonni-corrected) and reverse (p=0.0099; Bonferonni-corrected) replay rates for the trained sequence relative to this control (Fig. 2D; see also Fig. S2C for additional control sequences). Thus, replay during skill learning is selective for, but not exclusive to, the trained motor skill sequence.

### Replay of the learned skill is represented within a mediotemporal-sensorimotor network

We also examined the spatiotemporal features of replay event dynamics. After averaging all 50ms duration forward replay event epochs detected by the decoding pipeline during inter-practice rest, we applied principal component analysis (PCA) to extract low-dimensional networks with co-varying source-space power changes during fast replay. Two orthogonal networks explained 86.6% of the total signal variance during an average replay event. The first network (PC1 explaining 67.8% of total variance) featured sensorimotor, entorhinal, and hippocampal regions (Fig. 3). It also included the precuneus, one of the main outputs of the hippocampus to the neocortex^22, 23^, shown to be active in previous fMRI work during rest periods of early learning^9^. The second network (PC2) included a similar set of regions except for the precuneus (Fig. S4). Thus, replay of the skill is represented predominantly in sensorimotor, entorhinal and hippocampal regions.

**Figure 3.**
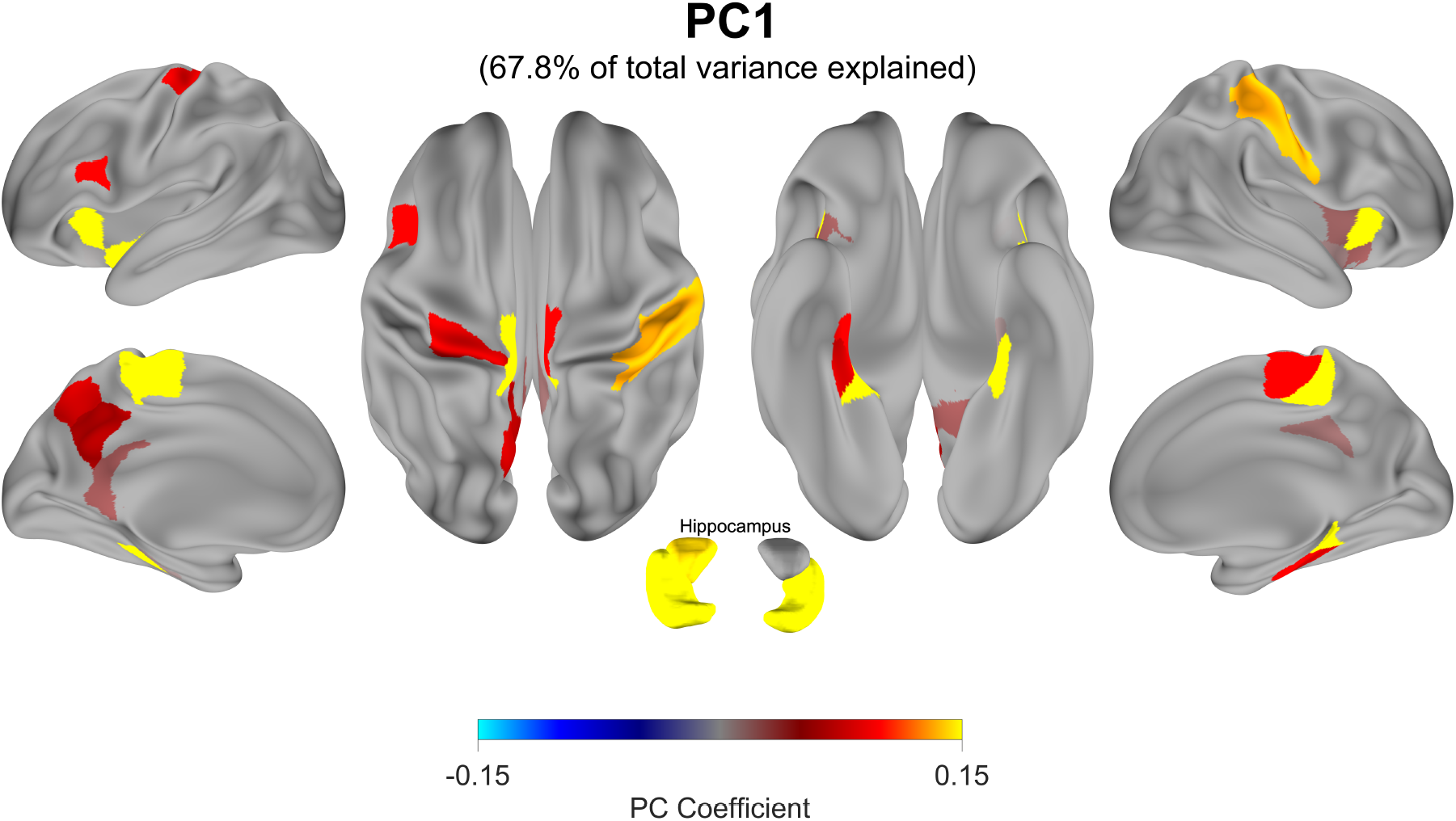
Spatial features of detected replay events. Principal component analysis (PCA) was used to characterize networks with covarying power changes in parcellated MEG source-space during group average replay events. Together, PC1 and PC2 explain 86.6% of the total power-related variance during replay. PC1, shown here, explains 67.8% of the total variance alone and is characterized by a positively correlated sensorimotor-entorhinal-hippocampal network. PC2 displays a similar set of regions (Fig**. S4).** Surface maps are thresholded to display the highest loading parcels (see Methods).

### Forward and reverse replay correlate with wakeful consolidation of skill

Finally, we asked if wakeful consolidation in individual participants could be predicted by the rate of replay events detected for the trained sequence during inter-practice rest intervals. We observed significant correlations between inter-practice neural replay rate and cumulative micro-offline gains (Fig. 4) for both forward (Fig. 4A; r=0.451, p=0.012) and reverse (Fig. 4B; r=0.521, p=0.003) replay of the trained sequence. On the other hand, replay of the control sequence (*33433*; see Methods) did not correlate with behavior (r=0.165, p=0.38). In summary, greater replay rates related to the trained sequence predicted greater rapid wakeful skill consolidation across individuals.

**Figure 4.**
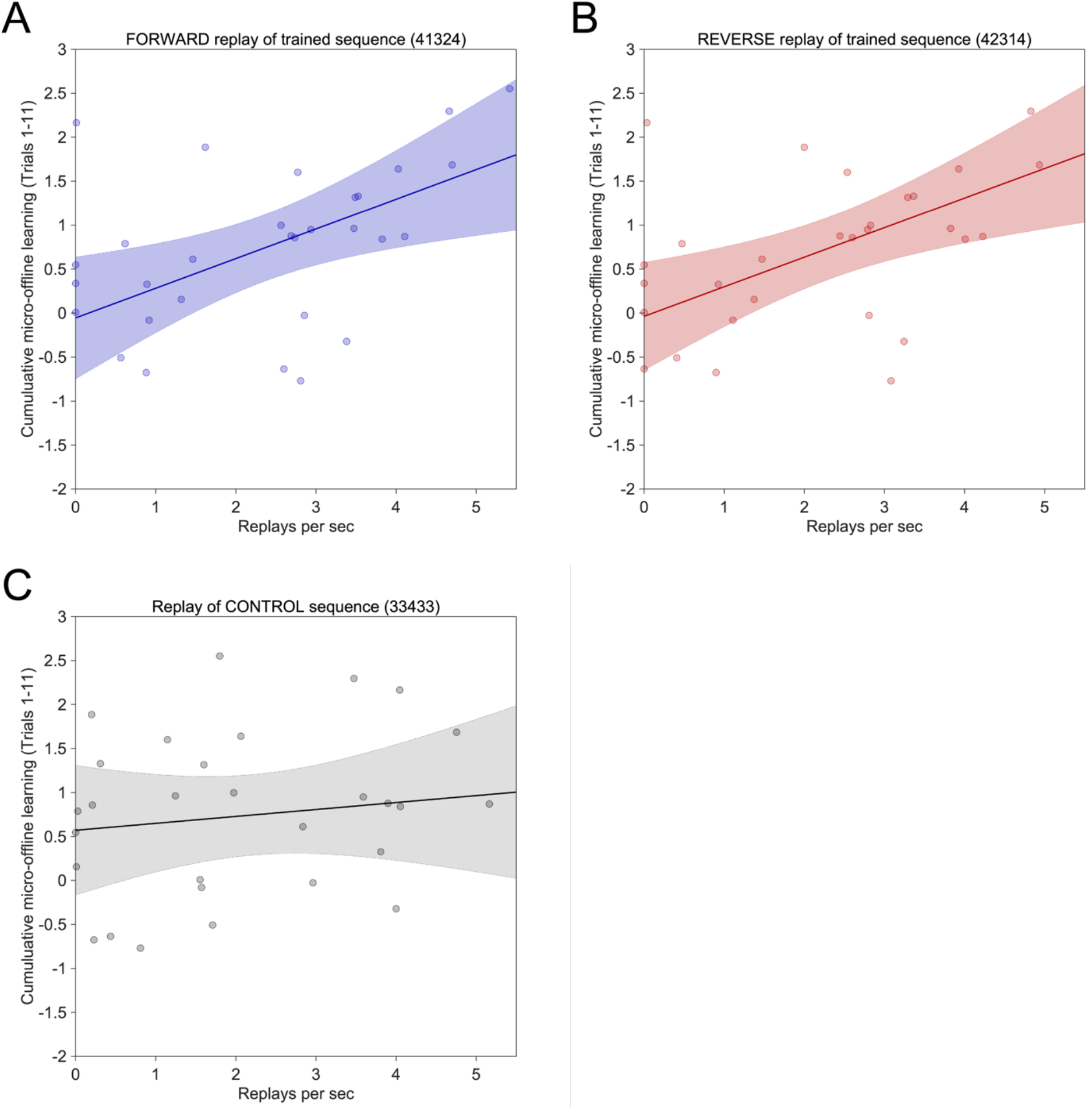
Forward and reverse neural replay of the trained sequence predict rapid wakeful skill consolidation. The above scatter plots depict inter-individual replay rates over all early learning inter-practice rest periods (abscissa) relative to cumulative micro-offline gains (ordinate) over the same time window (Trials 1-11). **(A,B)** Significant correlations were observed for both forward (A; r=0.451, p=0.012) and reverse (B; r=0.521, p=0.003) replay of the trained sequence. Thus, higher rates of replay related to the trained sequence predicted greater micro-offline gains and **rapid** wakeful skill consolidation in individuals. (**C)** Conversely, replay during inter-practice rest of the control sequence (*33433*) did not display a significant correlation with **rapid** wakeful skill consolidation (r=0.165, p=0.38) during early learning.

## Discussion

In this study, we address the question of how the brain consolidates memory during waking rest. Positing a role for neural replay, we first document replay in medial temporal and sensorimotor brain regions during wakeful human memory consolidation. We then demonstrate that replay is temporally compressed by approximately 20x relative to the practiced behavior. Finally, we show that forward and reverse waking replay rates increase rapidly after the onset of practice, display selectivity for the trained sequence, and predict consolidation of skill.

The presence of waking rest interspersed with practice influences successful learning. For example, skill memory is enhanced when practice events are separated by rest rather than massed in immediate succession, a phenomena known as the “spacing effect”. This observation, initially made by Ebbington in 1885^24^, has been repeatedly documented across cognitive domains^2, 3, 4, 5, 6^. Waking rest also contributes to skill acquisition. Early learning (a time period where performance improves exponentially) is largely accounted for by *micro-offline* gains that occur between, rather than during practice periods^8^. Thus, a form of wakeful consolidation of skill develops over seconds and minutes – much shorter timescales than previously thought^10^. Importantly, this form of wakeful memory consolidation is approximately four times greater in magnitude than classically studied overnight consolidation, which requires sleep^8, 9^, and is preserved even when practice exposure is reduced by half^10^. Here, we studied how the brain binds, during waking rest, sequences of discrete motor actions learned during previous practice, leading to successful consolidation of skill^25^.

Waking replay, displaying robust temporal features, was decoded during the same rest periods in which rapid consolidation occurs. To our knowledge, this is the first demonstration of fast waking human neural replay linked to memory consolidation. Replay of full practiced sequences had a modal duration of approximately 50ms, representing a 20-fold temporal compression of keypress sequence representations within each replay event relative to the duration of the actual skill behavior. This is consistent with the range of compressions previously reported in animal models as well as in humans^25, 26^, and is too fast to be explained by mental rehearsal or motor imagery processes which share a veridical duration with executed behavior (in this case, 1000-2500ms replay durations)^27, 28^.

Immediately following the initiation of practice, replay rates during interspersed rest periods increased more than three-fold beginning with the first one (Fig. S3A). This finding is reminiscent of the increase in waking replay rates in rodents upon introduction into a novel maze environment^15, 29^. We also found a high likelihood for temporal clustering, as more than half of all detected replay events were paired with a second event within a window of <200ms (Fig. S3D). Such replay clustering^25^ has been linked to memory consolidation^15^. Following practice, replay rates receded to pre-training levels rather rapidly contrary to the slow decrease reported in rodents^30^ – a difference that could be attributed to the explicit awareness of training termination in humans or to the presence of reward contingencies in rodents, absent in each of the other species^19^.

The increase in both forward and reverse waking replay^18^ rates for the trained sequence was larger than for the alternative unpracticed sequences examined (Fig. 2D, S2C). While forward replay of this trained sequence could support retrieval of information obtained during previous practice relevant for planning and guiding action in the next practice period^17, 30, 31^, reverse replay could contribute to the evaluation of recent practice outcomes.^17, 18, 19, 25^ Alternatively, the increase in reverse replay could have been driven by shared ordinal locations with the trained sequence. The fictive replay we observed for unpracticed sequences is not suprising given that studies of navigation memory in rodents have repeatedly shown that hippocampal cell assemblies generate plausible trajectories or sequences that have not yet been explored^32, 33, 34^. It is possible that such fictive replay events reflect more generalizable abstract knowledge structures (i.e. – cognitive maps) important for skill transfer^35^. The keypress sequence representations we also observed during pre-training rest after subjects had received task instructions but before they initiated practice (Fig. 1A, 2B) can be similarly explained. Evidence of *“pre-play”* in rodents indicates these reactivation patterns could be generated from intrinsic neural dynamics supporting broad sequence representations that are subsequently tuned through experience^32, 36^. Alternatively, it is also possible that semantic knowledge about the task requirements was enough to elicit replay of keypress sequences in preparation of the impending training ^25^.

Learning this skill requires binding individual discrete keypress actions into a fast, precise spatiotemporal sequence^37^. While neural representations of individual keypresses – stereotypical, overlearned finger movements – are highly invariant and maintained within the primary motor cortex^38^, keypress sequence skills are spatially represented throughout a distributed frontoparietal brain network that reorganizes with learning^9, 16, 37, 38^. Waking neural replay events characterized here were **encoded in regions that differed from those r**eported for practice sequences^38^ in the proportion of hippocampal-entorhinal engagement^11^. The most parsimonious explanation for this difference is that during the rest periods within which consolidation occurred, synchronization of hippocampus with engaged neocortical regions potentiated the skill memory to a greater extent than during practice itself^39, 40^. For individual replay events, which were detected on average between 20-30 times during each 10-second long rest period, the hippocampus and entorhinal cortex could engage memories encoding abstract knowledge about the action structure of the skill sequence, while the sensorimotor cortex may engage memories containing information about the kinematics and dynamics of the elemental finger movements. In this way, replay initiated by the hippocampus could promote the binding of abstract knowledge about the action structure of the skill with detailed sensory predictions important for execution and evaluation of the skill^41, 42, 43^. This view is consistent with evidence of hippocampal involvement during rest periods of early learning^9, 16^. Taken together, our data indicate that frequent, fast waking replay reinforces hippocampus **and** neocortical associations learned during prior practice, a process relevant for improving subsequent performance and wakeful consolidation of skill. Still, while the correlation between replay of the trained sequence and micro-offline learning is strongly suggestive of a direct contribution to consolidation of skill^8^, causality remains to be established in either animals or humans^44, 45^.

We conclude that robust hippocampal and neocortical replay predicts micro-offline learning and is a possible binding mechanism supporting wakeful memory consolidation.

## STAR ⋆ Methods

**Key Resources Table.**

| Reagent or Resource | Source | Identifier |
| --- | --- | --- |
| **Deposited Data** |  |  |
| MEG Data | This paper | N/A |
| **Software and Algorithms** |  |  |
| Matlab | Matlab | https://www.mathworks.com |
| Python | Python | https://www.python.org |
| E-prime 2 | Psychology Software Tools, Inc., Sharpsburg, PA, USA | https://pstnet.com/products/e-prime/ |
| FreeSurfer |  | https://surfer.nmr.mgh.harvard.edu/ |
| MNE |  | https://mne.tools/stable/index.html |
| Custom code and algorithms | This paper | https://github.com/Claudino/replay |
| **Other** |  |  |
| Human healthy volunteers recruited from local area | This paper | N/A |
| CTF MEG 275 system | CTF MEG International Services LP. Coquitlam, BC, Canada | https://www.ctf.com |
| Cedrus LS-LINE | Cedrus Corporation. San Paolo, CA, USA | https://www.cedrus.com/lumina/ |
| GE Discovery MR750 3T scanner system | GE Healthcare. Chicago, IL, USA | https://www.gehealthcare.com/products/magnetic-resonance-imaging/3-0t/discovery-mr750 |
| Siemens Skyra | Siemens Healthineers. Erlangen, Germany | https://www.siemens-healthineers.com/magnetic-resonance-imaging/3t-mri-scanner/magnetom-skyra |
| NIH HPC Biowulf Cluster | NIH, Bethesda, MD, USA | https://hpc.nih.gov/ |

### Resource Availability

#### Lead Contact

Further information and requests for resources should be directed to and will be fulfilled by the Lead Contact, Ethan R Buch.

#### Materials Availability

This study did not generate new materials.

#### Data and Code Availability

Data (subject to participant consent) and analysis code used to generate the findings of this study will be made freely available upon request to the Lead Contact, Ethan R Buch.

### Experimental Model and Subject Details

#### Participants

Thirty-three naïve right-handed healthy participants (16 female, mean ± SEM age 26.6 ± 0.87) with a normal neurological examination gave their written informed consent to participate in the project, which was approved by the Combined Neuroscience Institutional Review Board of the National Institutes of Health (NIH). Active musicians were excluded from the study ^46^. The sample size was determined *a priori* via a power analysis of prior skill learning data collected in our research group using the same task^47, 48^. Two participants were excluded before the start of analysis because of technical problems with the MEG scanner during the recording sessions. One additional participant was excluded due to large movement artefacts in the MRI scan required for source modeling.

## Methods

### Behavior

#### Task

Participants performed a novel explicit motor skill sequence task where they repetitively typed a 5-item numerical sequence displayed on a computer screen (*41324*) as quickly and as accurately as possible. Keypresses were performed with the participant’s non-dominant, left hand and applied to a response pad (Cedrus LS-LINE, Cedrus Corp). Keypress 1 was performed with the little finger, keypress 2 with the ring finger, keypress 3 with the middle finger and keypress 4 with the index finger. The thumb was not used to respond. Individual keypress times and identities were recorded for behavioral data analysis. Participants practiced the task for thirty-six (36), 10-second duration trials. 10-second rest periods were interleaved between trials. During practice, participants were instructed to fixate on the five-item sequence displayed for the full duration of the trial. Small asterisks appeared above a sequence item when a keypress was recorded, providing feedback to the participant about their current location within the sequence. After completion of a full iteration of the sequence, the asterisks were removed. The asterisk display did not provide error feedback, since they appeared for both correct and incorrect keypresses. During the 10-second interleaved rest periods, the sequence was replaced with a string of five “X” symbols, which participants were instructed to fixate on. Visual stimuli and task instructions were presented, and keypress responses recorded using a custom script running in E-Prime 2 (Psychology Software Tools, Inc.).

#### Skill measure calculation

Instantaneous correct sequence speed (skill measure) was quantified as the inverse of time (in seconds) required to complete a single iteration of a correctly generated full 5-item sequence. Individual keypress responses were labeled as members of correct sequences if they occurred within a 5-item response pattern matching any possible circular shifts of the 5-item sequence displayed on the monitor (*41324*). This approach allowed us to quantify a measure of skill within each practice trial at the resolution of individual keypresses and identify all possible types of keypress response errors (i.e. – omission, replication or substitution). Performance within each trial was summarized as the median instantaneous correct sequence speed over the full trial.

#### Determination of the early learning trial cutoff

Here, early learning corresponds to the set of trials between trial 1 and a cutoff trial, *T*, on which 95% of the performance gains are first achieved *at the group level*. We determined *T* as follows. First, we fit the trial-by-trial group average correct sequence speed data with an exponential model of the form:

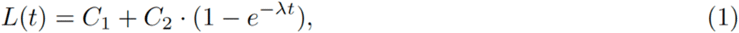

where *L(t)* represents the group average learning state on practice trial, *t*, *C_1_* and *C_2_* control the pre-training performance and asymptote (i.e. – extent of learning), and λ controls the learning rate. Parameters *C_1_* (boundry constraints = [0,5]), *C_2_* ([0,15]) and λ ([0,2]) were estimated using a constrained nonlinear least squares method (MATLAB’s *lsqcurvefit*, trust-region-reflective algorithm). Next, we accumulated *L(t)* (Eq. 1) over all $t$ and divided each trial’s cumulative sum by the area under the curve to obtain a trial-by-trial cumulative percentage of learning. Finally we set *T* to the first trial where 95% of the learning had been achieved (*T* = 11 in this study).

#### Calculation of micro-online and micro-offline gains during early learning

We then assessed early learning changes at the individual trial level by quantifying performance improvements occurring during individual practice and rest periods. *Micro-online* learning was defined as the difference in instantaneous correct sequence speed between the beginning and end of a practice trial. *Micro-offline* learning was defined as the absolute difference in correct sequence speed between the end of a practice period and the beginning of the subsequent practice period. Cumulative *micro-online* and *micro-offline* learning scores measured over the first 11 trials for each participant were used to assess their respective contribution to total early learning (i.e. - change in performance between Trial 1 and Trial 11)^8^.

### Magnetic resonance imaging (MRI)

#### MRI Acquisition

Structural MRI scanning was performed on a 3T MRI scanner (GE Excite HDxt and Siemens Skyra) with a standard 32-channel head coil. T1-weighted high-resolution (1mm^3^ isotropic MPRAGE sequence) anatomical images were acquired for each participant.

### Magnetoencephalography (MEG)

#### MEG Acquisition

Continuous MEG was recorded at a sampling frequency of 600Hz with a CTF 275 MEG system (CTF Systems, Inc., Canada) while participants were seated in an upright position. The system is composed of a whole head array of 275 radial 1^st^-order gradiometer/SQUID channels housed in a magnetically shielded room (Vacuumschmelze, Germany). Two of the gradiometers were malfunctioning and were not used. A third channel was removed (MLT16-1609) after visual inspection of the data indicated the presence of high noise levels for all recordings, resulting in 272 total channels of MEG data. Synthetic 3rd order gradient balancing was used to remove background noise online. A TTL trigger sent from the task computer to the MEG data acquisition computer was used to temporally align the behavioral and MEG data. Head position indicator (HPI) points were monitored in the scanner coordinate space using head localization coils (HLCs) placed on the nasion, left and right pre-auricular locations of the participant’s head. These coil positions were also recorded in the subject’s T1-MRI coordinate space during the session using a stereotactic neuronavigation system (BrainSight, Rogue Research Inc.).

#### MEG data preparation and pre-processing

Each participant’s MEG data were organized into the following samples: *pre-training rest* (single segment of 5 min), *inter-trial practice rest* (36 segments of 10s), *practice* (36 segments of 10s) and *post-training rest* (single segment of 5 min). All MEG samples were band-pass filtered between 1-100Hz, notch-filtered at 60Hz and subjected to Independent Component Analysis (ICA). ICA was fit using *MNE Python*’s^49^ implementation of *FastICA*^50^ after pre-whitening with Principal Component Analysis (set to retain the top 15 components). Spatial topographies and time series of all components were visually inspected and those determined to be strongly associated with known electrocardiogram, eye movement and blinks, and head movement artefacts were removed from the MEG sample.

#### MEG source reconstruction and parcellation

Source reconstruction of the MEG data was performed using the standard pipeline in MNE Python^51^. A single forward solution and inverse solution was calculated for each MEG sample. A boundary element model (BEM) required for generation of the forward solution was constructed for individual subjects from inner-skull and pial layer surfaces obtained by segmentation of the subject’s T1-MRI volume in *FreeSurfer*^52^. Hippocampal surface labels were obtained by parcellating the participant’s MRI according to the coordinates in the *Brainnetome* atlas^50^ (https://atlas.brainnetome.org/). Neocortical and hippocampal source spaces were obtained by sampling the corresponding surfaces on a recursive octahedral grid with 4.9mm between dipoles, and spatially aligned to the MEG sensor data with the co-registration data obtained from BrainSight. The BEM was then used to generate the forward solution at each source dipole location. The Linearly Constrained Minimum-Variance (LCMV) beamformer operator was used to computer the inverse solution from the recorded MEG sensor space data. The entire duration of each MEG sample was used to calculate the inverse solution data covariance matrix. All inverse solutions used the same sample noise covariance matrix, which was calculated over the first 20s of the pre-training rest MEG sample for individual subjects.

MNI-space transformations obtained from the FreeSurfer segmentation pipeline were then used to spatially register the resulting source reconstruction time series to the *Brainnetome* atlas, enabling quantitative estimation of millisecond-resolved brain activity for 212 neocortical and 4 hippocampal parcels (i.e. – 216 total parcels). Parcel activity was estimated by averaging the time series of all source dipoles falling within a given parcel boundary. A process called *mean-flipping* was used to prevent within-parcel source cancellation when averaging. That is, the sign was flipped for all sources with a sign that differed from that of the average source. Since the sensor space data had been bandpass filtered with a cut-off frequency of 100Hz, the source data was downsampled from 600Hz to 200Hz to increase computation speed in subsequent processing steps. All remaining analyses were performed directly on the parcellated source-space MEG time series.

#### Training and evaluation of keypress decoders

Individualized keypress decoders were trained for each participant using Python’s *Scikit-learn* library^53^. The training data of a subject’s decoder was assembled as follows. We time-averaged the parcellated source for each correct keypress in all 36 practice MEG samples between *t* ± *Δt* ms, where *t* is the keypress timestamp and *Δt* is treated as a cross-validated hyperparameter described in detail below. The timestamp for each keypress was obtained from behavioral data files output by E-prime. This produced the predictor matrix **X**_[*keypresses* × *parcels*]_ and the target array ****y****_[*keypresses* × 1]_ where *keypresses* varied between subjects (some typed more sequences than others) and parcels=216 as earlier described.

We used **X** and ****y**** to train four one-versus-all Radial-basis Support Vector Machines (Python *Scikit-learn’s* SVC) with a pipeline parameterized with a feature centering/scaling policy, a time window for averaging the MEG source around a keypress (*Δt*), a regularization term (*C*) and a kernel coefficient (*γ*). We solved for these (hyper)parameters by running 5-fold cross-validated grid-search hyperparameter optimization three times, one for each of the following optimality criteria:

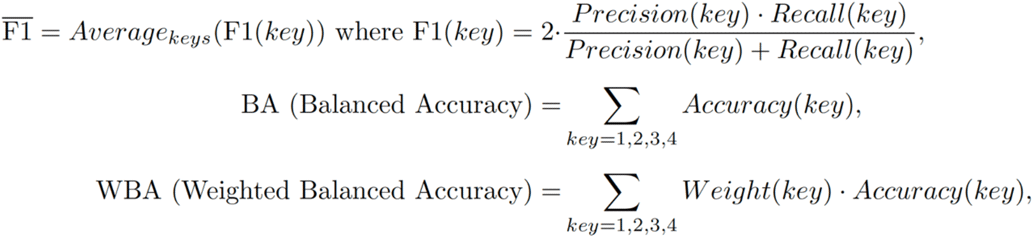

where *Weight*(*key*) = [2/7, 2/7, 2/7, 1/7] for keys 1, 2, 3 and 4, respectively, and *Accuracy*(*key*) = percentage correct classification of *key*. For the second and third optimality criteria, classification was done by picking the *key* that maximized the probability estimated by the SVCs, rather than using the decision function. The search grid used was: *C* = [0.1, 0.5, 1, 10, 50, 100, 1000], *gamma*=[0.05, 1/(2 × *keypresses*), 0.1, 0.5, 1], *center policy* = [True (will remove median), False (will not not remove median)], standardization policy = [True (will divide by inter-quartile range (IQR)), False (will not divide by IQR)] and *Δt* =[5, 10, 25] – that is a total of 420 configurations. SVCs were trained with a target error of 10^−3^ or less.

Three cross-validated grid searches were run for each participant, one for each of the three criteria : 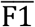, BA and WBA. Grid search under a particular criterion worked as follows. First, the training set was divided into 5 mutually exclusive stratified partitions so that the proportions of all keypresses were preserved as in the original data (since the training sequence was *41324*, each folder would have approximately twice as many *keys*=4). Next, we used the random oversampler in Python’s *Imbalanced-learn* toolbox^54^ (with a “not majority” sampling policy) to independently oversamples keypresses 1, 2 and 3 within each training partition. Note that test partitions were never oversampled, and training partitions did not share common oversampled patterns as this would lead to double-dipping and ultimately overfitting. Finally, we performed the same 5-fold cross-validation procedure on each of the 420 configurations in the grid. Each configuration produced 5 estimates, resulting in a total of 2100 estimates. We then selected the decoder configuration with the best cross-validated performance (data in test partitions only, and performance under the considered optimality criterion) for each individual participant. To evaluate the decoder’s performance, we created a null performance set by permuting the test labels (i.e. – identities of all keypresses) 1000 times and calculating the decoder’s performance after each permutation, leading to a *p-value* = (*correct classifications* + 1) / (1000 + 1). We also calculated the decoder’s cross-validated confusion matrix, that is, the confusion matrix calculated over the data in the cross-validated test partitions. This confusion matrix played an important role in our replay detection pipeline as described later.

#### Estimation of decoder performance on practice sequence detection

The goal of the present step was to select the best decoder for a participant on the basis of their estimated performance on full practiced sequences. Recall that keypress decoder training resulted in three decoder options for each participant (i.e. –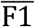, BA and WBA) with associated cross-validated performances, p-values and cross-validated confusion matrices. We used the following process to select the optimal decoder for each participant. First, we discarded any decoder with a *p-value* >= 0.05. This step resulted in at least one viable decoder for all participants. Next, we reshaped each participants’ predictor matrix **X**^Practice^_[*keypresses* × *parcels*]_ back into the correctly typed sequences they originated from, resulting in an associated array **X**^Practice^_[*5* × *parcels* × *N*]_ for each subject, where *N* is the number of correctly typed sequences of 4-1-3-2-4.

We then evaluated the ability of the decoders to correctly identify each typed practice sequence from *j*=1…*N* using the following procedure:

1. Scored the probability, *p_seq_*[*j*], of **X**^Practice^_[:, :, *j*]_ being identified as sequence 4-1-3-2-4 as the product of the probabilities of each of the sequence’s constituent keys, *p_key_*(⋅), in the correct order. Probabilities *p_key_*(⋅) were obtained from the keypress SVC decoders for each subject.
2. Repeated Step 1 for all other possible sequences (i.e. – all other 1023 5-digit combinations of 1,2,3 and 4). Combined, these sequences form the set of null probabilities, Ω.
3. From Ω, we calculated the p-value for detection of the *j-*th practiced sequences as p-value = 1 − *percentile*^Ω^(*p_seq_*[*j*])/100 and counted a detection of typed sequence *j* if the p-value < 0.05.

Finally, we obtained the sequence detection rate of a decoder by dividing the number of detections by the total number of practiced sequences, *N*. We then selected the decoder option with the top detection rate for sequences observed in the practice data. Fig. S5A shows the distribution of top detection rates across participants. Fig. S5B shows keypress probabilities first averaged at each ordinal position of detected practice sequences and then averaged over subjects. Note that the expected key at each position is always the keypress with the top detection probability. All participants had at least one decoder with a non-zero detection rate. The selected decoder was the one used to interrogate waking rest MEG data samples for replay events.

#### Sequence replay detection and replay rate calculation

Given a rest period MEG sample, **X** ^Rest^ _[*T* × *parcels*]_, of *T* time-points *x* parcels recorded from a given subject, we estimated the probability that a given sequence, *seq*=ABCDE, was replayed at a time 0 ≤ *t* < *T* within that sample to be the product of the probabilities of each of the sequence’s constituent keys A, B, C, D and E ∈ [1, 2, 3, 4] being observed both in the right order and ending at time *t*, that is:

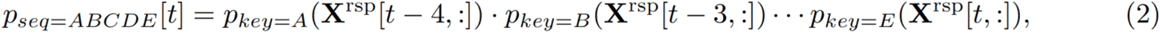

Where **X**^rsp^ = resample(**X**^Rest^[t − 5 · tscl + 1 : *t*, : ]) is an MEG sample segment extracted from t − 5 · tscl + 1 to *t* at all 216 parcels and compressed down to 5 time-points. Probabilities of different replay durations were obtained by varying the timescale compression factor, *tscl*, where *tscl* ∈ [1, …,10,15,20,30,40,80,100].

For example, since our time resolution is 5ms, we set *tscl* = 1 to look for 25ms replay durations, *tscl* = 2 for 50ms replay durations, and up to as high as *tscl* = 100 for 2500ms durations. In terms of implementation, we obtained **X**^*rsp*^ using Python’s *scipy resample* with default arguments and *num*=5. Again, probabilities *p_key_*(·) were obtained from the keypress SVC decoders selected for each subject. *Scikit-learn* implements a multiclass version of Platt’s scaling technique to obtain probability distributions from classifications over more than two classes^55^.

We only assessed replay likelihood at time-points where *p_seq=ABCDE_*[*t*] exceeded a threshold, *θ_seq_*, set to the product of the maximum off-diagonal detection rates (i.e. false positive rates) of the keypress decoder’s cross-validated confusion matrix. For each time-point, *t*, such that *p_seq=ABCDE_*[*t*] > *θ_seq_*, we then tested the associated statistical significance as to “how rare” *p_seq=ABCDE_*[*t*] would be under the null distribution of probabilities for all other 1,023 possible decodable sequences (i.e. – the Ω set) resulting in a p-value = 1 − *percentile*^Ω^(*p_seq_*[*t*])/100. All p-values were FDR-corrected (Benjamin-Hochberg) to account for multiple comparisons using an α = 0:05 with the number of tests set to the size of Ω. By pre-selecting which time-points to test, we mitigated underpowering effects of correcting for multiples comparisons. Note that the selection threshold *θ_seq_* was fully determined from practice data (cross-validated confusion matrices) and never from test data on which statistical testing was carried out, therefore eliminating the possibility of circular analysis or double-dipping. Statistically significant replay detections were marked for all p-value[*t*] < 0.05. Since it was possible that overlapping X^rsp^[t] could result in multiple detections of the same putative replay event, each set of multiple detections occurring (within the same tested replay duration) was replaced with a single detection at the median time instant of the set. Finally, we scored the replay rate of *seq* during the given rest period MEG sample as the number of detections per the sample’s duration in seconds:

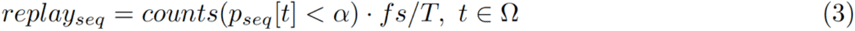

where *counts*(·) is the number of detections of *seq* along the timeline of the MEG sample, and *fs* is the sampling rate of the MEG sample (i.e. – fs/T is the inverse of the duration of the sample in seconds).

#### Replay Network Characterization

While use of a nonlinear kernel optimized the performance of the keypress state decoders, it precludes direct assessment of the spatial representations of these states (and sequence dynamics) from the decoder weights. However, since the output of the replay detection pipeline specifies the time-point at which individual replay events are initiated and what their duration is, it is possible to gain insight into replay network dynamics by interrogating these event epochs. After averaging all 50ms forward replay (i.e. . – 41324) event epochs, we performed principal component analysis (PCA) to extract low-dimensional networks displaying strong covariation in source power during replay. We mapped the membership coefficients for cortical regions onto the FreeSurfer average brain, and the four hippocampal subregions onto a separate surface representation. The thresholded maps display the top 10% of PC loading coefficient magnitudes (i.e – absolute value of signed coefficients).

#### Correlation of Replay Rates with Behavior

We performed Pearson correlation analyses of the average forward and reverse trained sequence replay rates detected during the first eleven 10-second waking rest periods with cumulative micro-offline learning over the same period. We also performed a control analysis by the assessing the correlation between replay of a single alternative sequence (*33433*) with with cumulative micro-offline learning over trials 1-11. This is the only alternative sequence (1 out of 1022) which does not share any common ordinal position or transition structure with the forward and reverse of the trained sequence.

## Acknowledgements

This research was supported by the Intramural Research Program of the NIH, NINDS. We thank Iñaki Ituratte for helpful comments on manuscript drafts.

## Author Contributions

E.R.B. conceived of the presented idea. M.B. and L.G.C. conceived of and designed the experiment. M.B. carried out the experiment. L.C. conceived of and developed the MEG analysis pipeline. R.Q. assisted with refinement of the MEG analysis pipeline. E.R.B. and L.C. carried out the analyses. E.R.B., L.G.C. and L.C. wrote the manuscript. M.B. and R.Q. reviewed and edited the manuscript.

## SUPPLEMENTARY FIGURES

**Figure S1.**
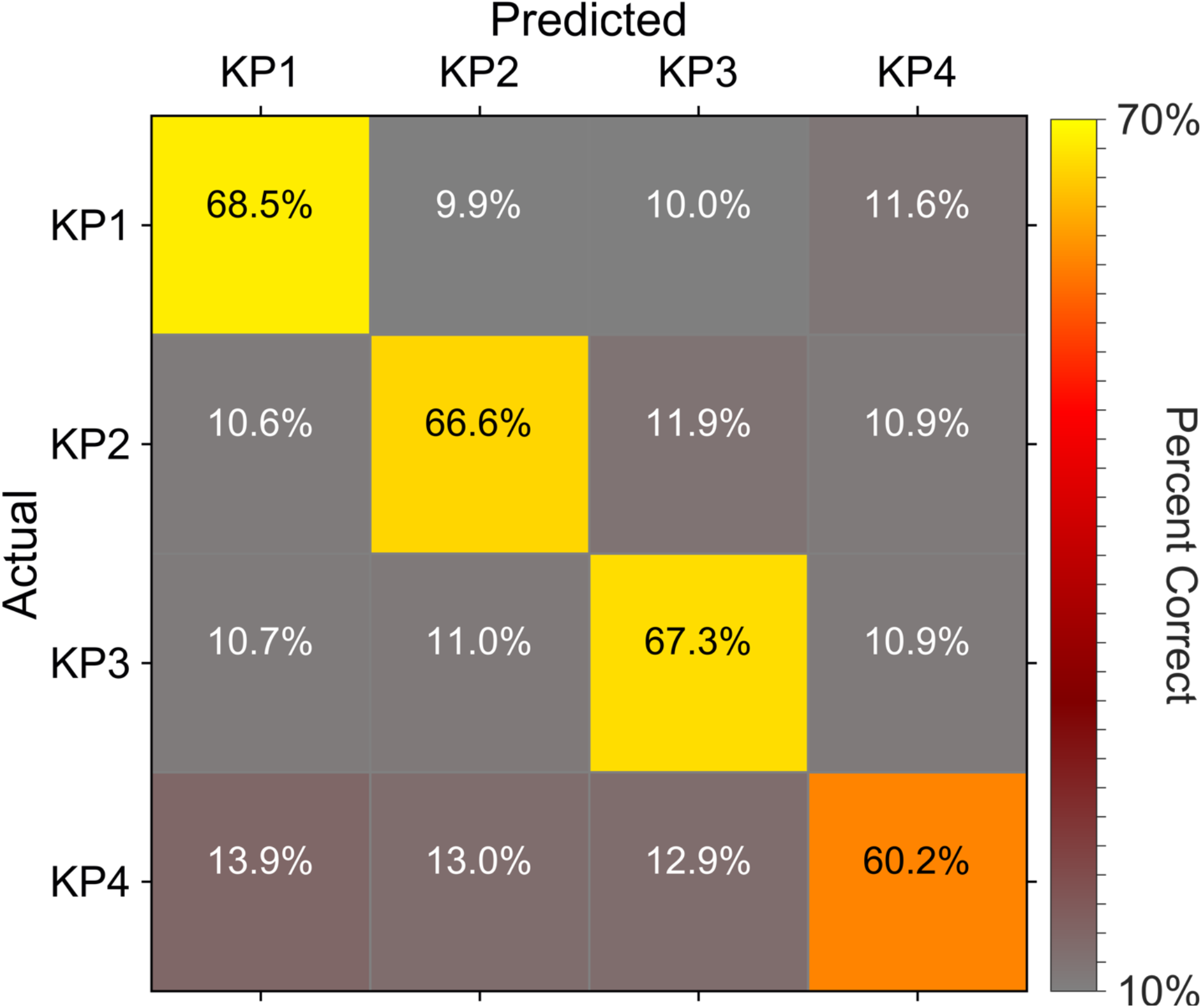
Subject-average confusion matrices of key press decoders calculated over training examples (average true positive rates ranged between 60.2-68.5%). Cross-validated performance of all selected decoders exceeded chance levels (Random permutation test, N=1000, α=0.05). Please see *Methods: Magnetoencephalography: Training and evaluation of keypress decoders* for complete details.

**Figure S2.**
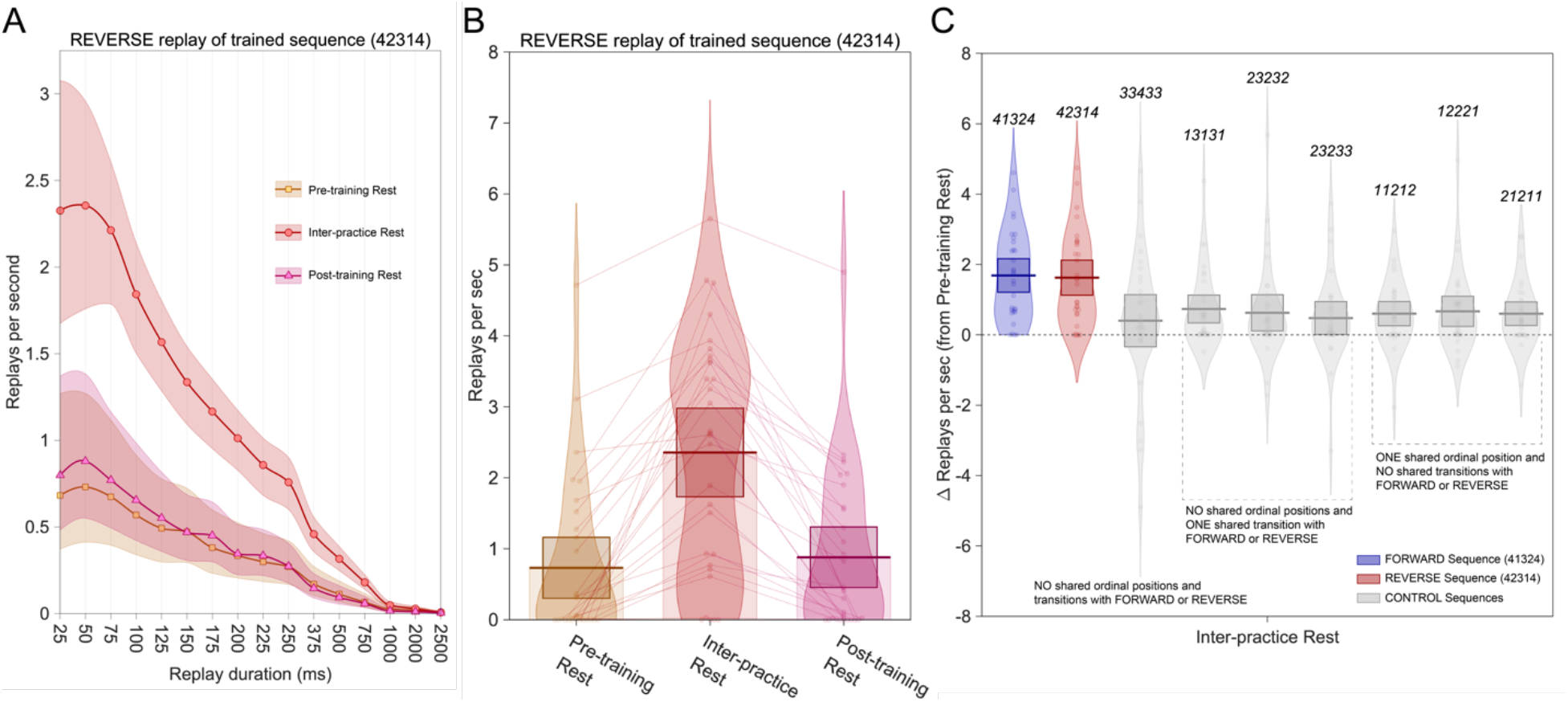
Reverse trained sequence replay detections. **(A)** The maximum reverse trained sequence replay rate was observed for events of 50ms duration. This was consistent with observed replay for the forward trained sequence (Fig. 2B). **(B)** Waking reverse replay rates (50ms-duration) for the forward trained sequence more than tripled during inter-practice rest (2.36±0.31) relative to pre-training rest (0.73±0.21). Similar magnitude rate changes were also observed for the forward trained sequence (Fig. 2C). Replay rates then receded during post-training rest (0.88±0.21) to levels similar to pre-training. **(C)** Forward and reverse replay of the trained sequence increased more prominently than all members of an expanded control sequence set after practice onset. Comparisons were limited to sequences sharing no common ordinal position or transition structure (n=1; *33433*), or only a single common ordinal position (n=3; 11212, 12221, 21211) or transition (n=3; 13131, 23232, 23233). Importantly, pre-training rest replay rates did not significantly differ between these sequences (F_8,108.48_=0.654; p 0.73).

**Figure S3.**
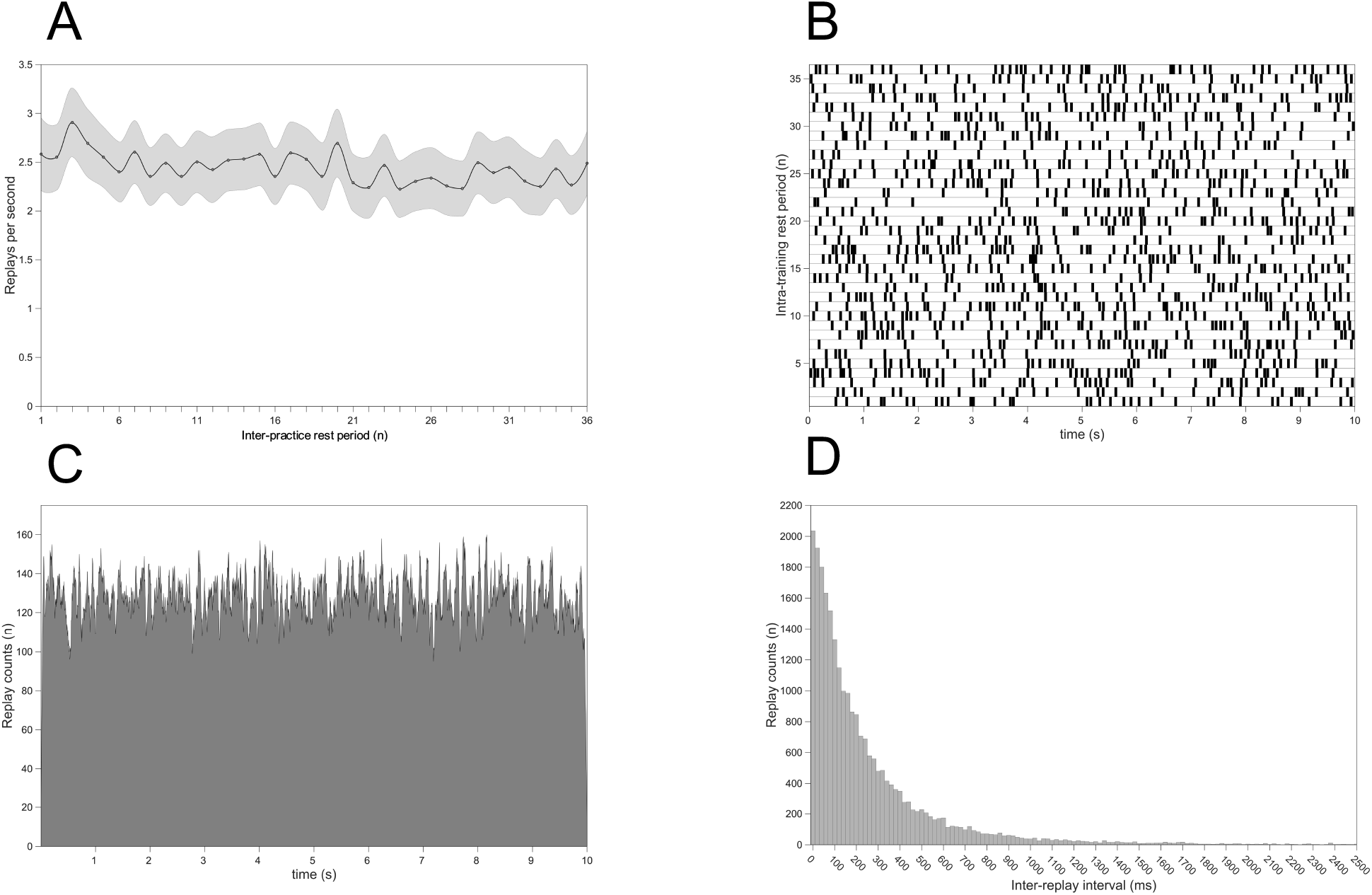
Temporal features of waking replay events during inter-practice rest. (A) High replay rates centered around 2.5/sec were detected over all 35 waking inter-practice rest periods. (B) Raster plot of inter-practice replay events (50ms duration) shown for a representative subject. (C) Histogram of replay event occurences summed across all 35 inter-practice periods for all subjects. Consistent with *(B)*, replays events were uniformly distributed across the entire 10-second rest interval. (D) Histogram showing the inter-replay intervals (i.e. – time between successive replay detections in ms) across all 50ms replay events in all subjects. The predominance of very short inter-replay intervals confirms that inter-practice rest replay events tend to occur in temporal clusters, consistent with the raster plot in *B*.

**Figure S4.**
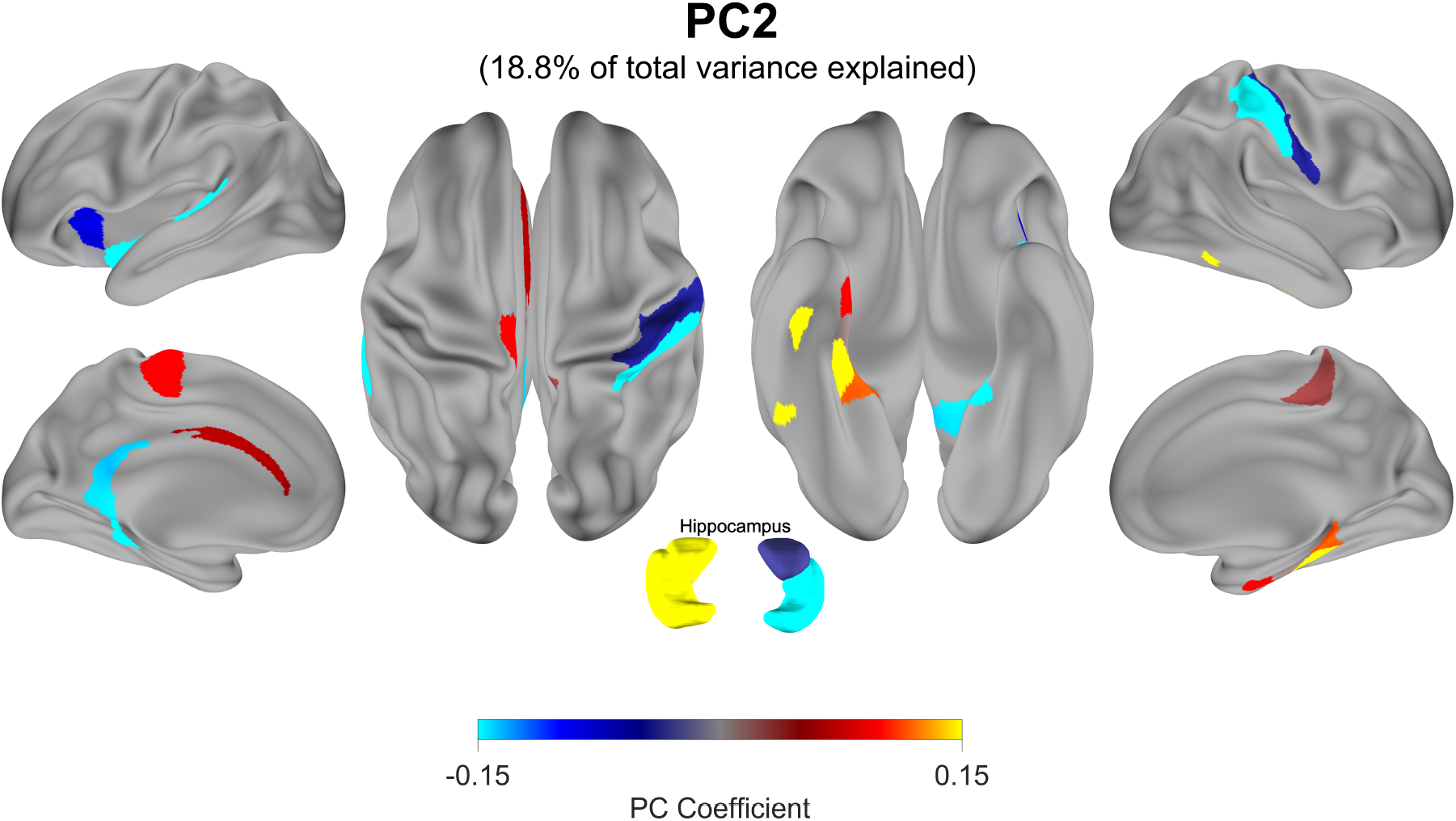
Spatial features of detected replay events. We performed principal component analysis (PCA) to characterize networks with covarying power changes in parcellated MEG source-space. PC2, shown here, explains 18.8% of the total power-related variance during group average replay events. Similar to PC1 (Fig. 3), PC2 and is characterized by strong covariations in power within a sensorimotor-entorhinal-hippocampal network.

**Figure S5.**
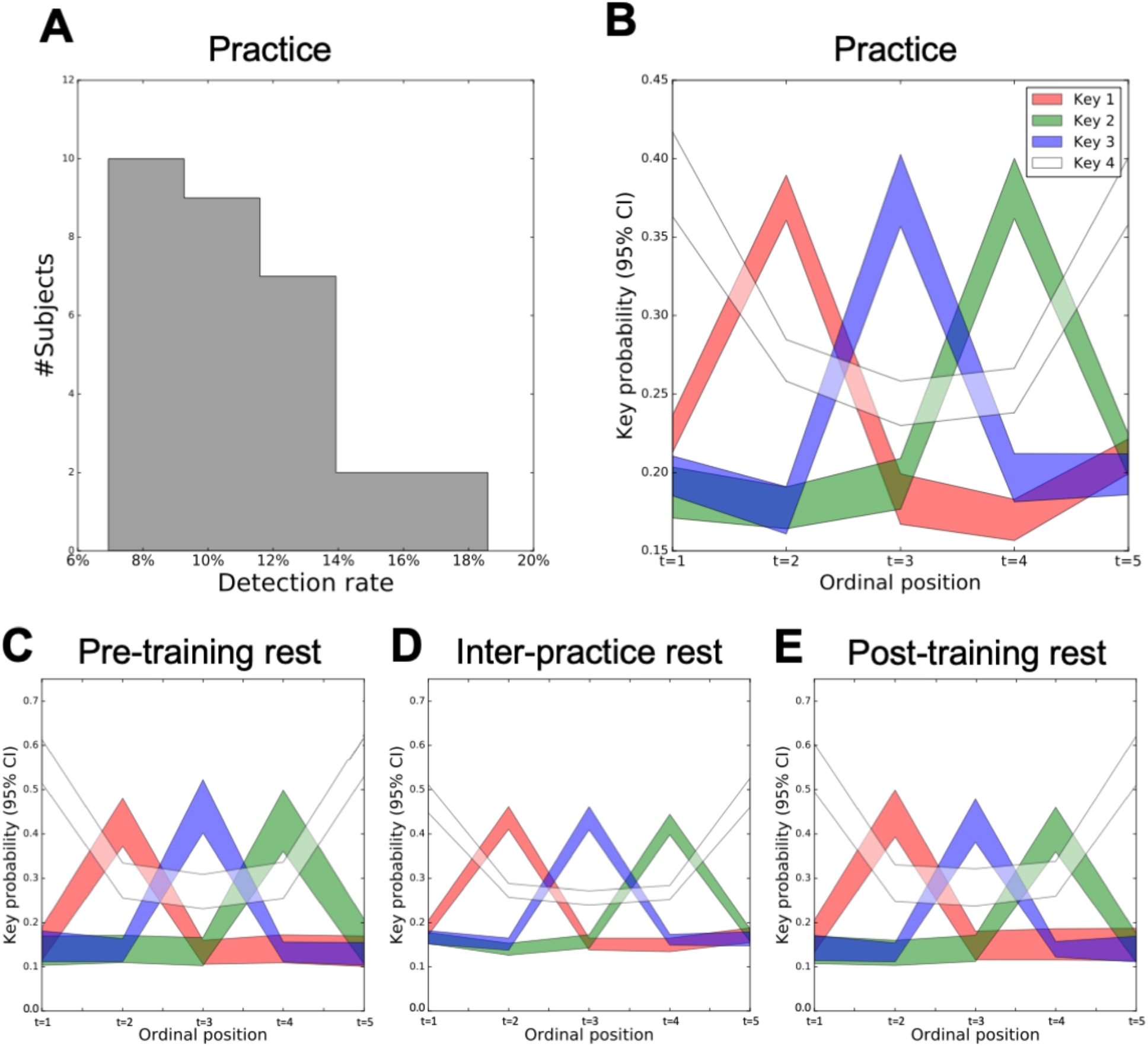
Estimation of decoder performance on practice sequence detection. **(A)** Distribution of sequence detection rates (percent of practice sequences correctly decoded from MEG activity only) over subjects. **(B)** Keypress state probabilities averaged at each ordinal position over detected practice sequences for all subjects (mean± SEM). Note that the maximum keypress state probability at each ordinal position correctly reconstructs the trained sequence (i.e. – *41324*). (**C, D, E)** Keypress state probabilities averaged at each ordinal position over all detected trained sequence replay events for all subjects (mean± SEM) during **(C)** Pre-training rest, **(D)** Inter-practice rest and **(E)** Post-training rest. Again, note that on average the maximum keypress state probability at each ordinal position is capable of decoding the full trained sequence (i.e. – *41324*) for replay events during waking rest.

